# Numb prevents a complete EMT by modulating Notch signalling

**DOI:** 10.1101/183871

**Authors:** Federico Bocci, Mohit Kumar Jolly, Satyendra C. Tripathi, Mitzi Aguilar, Samir M Hanash, Herbert Levine, José N. Onuchic

## Abstract

Epithelial-Mesenchymal Transition (EMT) plays key roles during embryonic development, wound healing, and cancer metastasis. Cells in a partial EMT or hybrid epithelial/mesenchymal (E/M) phenotype tend to exhibit collective cell migration, forming clusters of circulating tumour cells – the primary drivers of metastasis. Activation of cell-cell signalling pathways such as Notch fosters a partial or complete EMT, yet the mechanisms enabling cluster formation remain poorly understood. Using an integrated computational-experimental approach, we examine the role of Numb – an inhibitor of Notch intercellular signalling – in mediating EMT and clusters formation of hybrid E/M cells. Knockdown of Numb in stable hybrid E/M cells H1975 results in a full EMT, thereby showing that Numb acts as a brake for a full EMT. Consistent with this observation, we show via a mathematical model that Numb inhibits a full EMT by stabilizing a hybrid E/M phenotype. Thus, Numb can behave as a ‘phenotypic stability factor’ by modulating Notch-driven EMT. By generalizing the mathematical model to a multi-cell level, Numb is predicted to alter the balance of hybrid E/M versus mesenchymal cells in clusters, potentially resulting in a higher tumour-initiation ability. Finally, Numb correlates with a poor survival in multiple independent lung and ovarian cancer datasets, hence confirming its relationship with increased cancer aggressiveness.

Major Findings: we adopt an integrative computational-experimental approach to identify that Numb, an inhibitor of Notch signalling, can stabilize a hybrid epithelial/mesenchymal (E/M) phenotype. We show that knockdown of Numb in H1975 cells that display a stable hybrid E/M state is sufficient to destabilize a hybrid E/M state and push them to a full EMT phenotype. Next, we develop a mechanism-based mathematical model that recapitulates this ability of Numb in maintaining a hybrid E/M state, and predicts that Numb can alter the relative frequency of hybrid E/M and mesenchymal cells at a tissue level or in clusters of circulating tumor cells (CTCs) – the primary drivers of metastasis. Finally, we show that across cancer types, Numb correlates with worse patient survival, thus reinforcing the emerging notion that a hybrid E/M, but not necessarily a completely mesenchymal, phenotype associates with elevated tumour progression.

## Quick guide to equations and assumptions

The level of a generic protein, mRNA or micro-RNA (miR) X is described via a chemical rate equation that assumes the generic form:

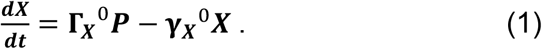

In (eq. 1), the first term on the right-hand side represents the production of X while the second term accounts for the degradation of X. Specifically, Γ_x_^0^ and γ_x_^0^ are basal production and degradation rates, while the functions P represents possible transcriptional/translational/post-translational regulations in the production rate of X and can be a function of any protein/mRNA/micro-RNA in the system.

Specifically, if the production rate of any chemical species X is increased/decreased in presence of an activator/inhibitor A, the overall production rate Γ_x_ = Γ_x_^0^ *P* takes the form:

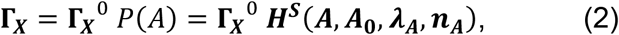

where *H^s^* is the shifted Hill function^12^, defined as:

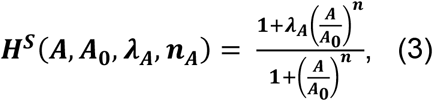

In (eq. 3), A is the level of activator/inhibitor expressed in number of molecules, *A_0_* is a threshold level, *n_A_* is the Hill coefficient and *λ_A_* represents the fold change in the production rate of X due to A. If A is an activator of X, then *λ_A_*>1 and the production rate is increased. Conversely, if A is an inhibitor of X, *λ_A_*<1 and the production rate is decreased.

## Introduction

Epithelial-Mesenchymal Transition (EMT) and its reverse Mesenchymal-Epithelial Transition (MET) play crucial roles during embryonic development, wound healing, and tumour progression^1^. Typically, cells that undergo EMT lose cell-cell adhesion and gain the traits of migration and invasion. These bidirectional transitions are rarely ‘all-or-none’. Instead, cells can display one or more hybrid phenotype(s) that possess a mix of epithelial and mesenchymal traits, thereby biasing them to undergo collective cell migration, instead of individual migration enabled by a complete EMT^1^. Collective migration, where cells maintain physical contact with their neighbours, has been considered to be a hallmark of multiple developmental processes such as neural crest migration, branching morphogenesis, and wound healing^1^. Recent studies have emphasized that collective cell migration can be a predominant path for cancer metastasis as well. Collective cell migration can enable the formation of clusters of Circulating Tumour Cells (CTCs)^2^. As compared to individually disseminating CTCs, these clusters are highly resistant to cell death in circulation, possess high tumour-initiation ability, and correlate with a worse clinical outcome across different cancer types^3^. Therefore, deciphering the intracellular and intercellular mechanisms that enables CTC clusters is essential to curb metastatic load.

The formation of clusters of CTC typically requires two conditions. First, individual cells can display a phenotype capable of both adhesion and migration, as is usually found in a hybrid epithelial /mesenchymal (E/M) phenotype. Further, such cells must be spatially co-located. It is theoretically possible that cells become hybrid in a random pattern and then dynamically find each other after the transition, but this is much more complex and hence less likely. Cellular mechanisms maintaining a hybrid E/M phenotype have received much attention^4-7^, particularly in a panel of non-small cell lung cancer (NSCLC) cell lines, but cell-cell communication in chemical and/or mechanical ways that can foster the direct formation of clusters via spatial organization remain relatively less studied.

Previously, we reported that Notch-Jagged signalling may both increase the frequency of cells in a hybrid E/M phenotype and their spatial proximity to form clusters of CTCs^8^. Notch signalling is an evolutionarily conserved cell-cell communication signalling pathway comprising a transmembrane receptor, Notch, and two transmembrane ligands, Delta and Jagged. When Notch binds to Delta or Jagged of a neighbouring cell, Notch is cleaved to release Notch Intra-Cellular Domain (NICD) that enters the nucleus, activates the Notch pathway and regulates its target genes^9^. NICD activates the transcription of Notch and Jagged, but represses that of Delta^10^. Thus, Notch-Jagged signalling between two neighbouring cells leads to convergent cell fates (lateral induction)^11,12^, whereas Notch-Delta signalling to divergent cell fates (lateral inhibition)^10^. Consequently, neighbouring hybrid E/M can reinforce stability of hybrid E/M phenotype and lead to formation of clusters of hybrid E/M cells via Notch-Jagged signalling^8^.

Based on this proposed role of Notch-Jagged signalling in inducing and maintaining a hybrid E/M phenotype, we hypothesized that the proteins affecting Notch signalling may modulate the stability of a hybrid E/M phenotype. Here, we focused on an inhibitor of Notch signalling – Numb^9^ – that promotes the degradation of Notch. Also, activated Notch signalling can inhibit Numb and its homolog Numb-like *(Numbl)*, generating a mutually inhibitory feedback loop between Numb and Notch^9^. Here, we first show that the knockdown of Numb in H1975 lung cancer cells that can maintain a stable hybrid E/M phenotype destabilizes the hybrid E/M state and pushes them towards a complete EMT. To rationalize these observations, we extended our mathematical model to allow for decoding the dynamics of the Numb-Notch-EMT signalling axis. We find that Numb can prevent the cells from undergoing a complete EMT, irrespective of the ligand (Delta or Jagged) that activates the Notch signalling. Thus, we demonstrate that Numb may behave as a ‘phenotypic stability factor’ (PSF) for a hybrid E/M phenotype.

Patient data indicates that Numb is correlated with poor survival. Under the assumption that clusters with high numbers of hybrid E/M cells have both the survival and tumorigenic properties needed to effectively seed metastases, we can address this issue with a spatially extended version of our mathematical model. Here we show that Numb can increase the percentage of hybrid E/M cells in clusters that undergo EMT, and is thereby consistent with the above finding. This last point can be addressed in future experiments which measure spatial patterns of Notch activity in both in *vitro* culture settings as well as *in vivo* CTC cluster analysis.

## Methods

### Cell culture and transfection

H1975 cells were obtained from the American Type Culture Collection. All cells grew in RPMI 1640 with 10% fetal bovine serum and a 1% penicillin/streptomycin cocktail (Thermo Fisher Scientific, Waltham, MA). Cell lines were cultured continuously for 6 months or less. Cell lines were validated at the MD Anderson Sequencing and Microarray Facility using short-tandem repeat DNA fingerprinting and routinely checked for mycoplasma by PCR (PromoKine). Cells were transfected at a final concentration of 50 nM siRNA using Lipofectamine RNAiMAX (Thermo Fisher Scientific) according to the manufacturer’s instructions using following siRNAs: siControl (Silencer Select Negative Control No. 1, Thermo Fisher Scientific), siNumb #1 (SASI_Hs01 _00245422, Sigma-Aldrich, St.Louis, MO), siNumb #2 (SASI_Hs02_00305013, Sigma-Aldrich, St.Louis, MO),), siNumb-L #1 (SASI_Hs01_00103891, Sigma-Aldrich), siNumb-L #2 (SASI_Hs01_00103892, Sigma-Aldrich). siRNA treatment were done for 48 hrs and then replated on coverslips for 24 hrs and then fixed.

### RT-PCR

Total RNA was isolated following manufacturer’s instructions using RNAeasy kit (Qiagen). cDNA was prepared using iScript gDNA clear cDNA synthesis kit (Bio-Rad). A TaqMan PCR assay was performed with a 7500 Fast Real-Time PCR System using TaqMan PCR master mix, commercially available primers, and FAM™-labeled probes for CDH1, VIM, ZEB1, NUMB, NUMBL, and JAG1 and VIC™-labeled probes for 18S, according to the manufacturer’s instructions (Life Technologies). Each sample was run in triplicate. Ct values for each gene were calculated and normalized to Ct values for 18S (ΔCt). The ΔΔCt values were then calculated by normalization to the ΔCt value for control.

### Western Blotting Analysis and Immunofluorescence

Cells were lysed in RIPA lysis assay buffer (Pierce) supplemented with protease and phosphatase inhibitor. The samples were separated on a 4–15% SDS-polyacrylamide gel (Biorad). After transfer to PVDF membrane, probing was carried out with primary antibodies and subsequent secondary antibodies. Primary antibodies were purchased from the following commercial sources: anti-CDH1 (1:1000; Cell Signaling Technology), anti-vimentin (1:1000; Cell Signaling Technology), anti-Zeb1 (1:1000; Cell Signaling Technology), anti-JAG1 (1:1000; Abcam), anti-Numb (1:1000; Abcam), anti-Numb-L (1:1000; Abcam) and anti-GAPDH (1:10,000; Abcam). Membranes were exposed using the ECL method (GE Healthcare) according to the manufacturer's instructions. For immunofluorescence, cells were fixed in 4% paraformaldehyde, permeabilized in 0.2% Triton X-100, and then stained with anti-CDH1 (1:100; Abcam) and anti-vimentin (1:100; Cell Signaling Technology). The primary antibodies were then detected with Alexa conjugated secondary antibodies (Life technologies). Nuclei were visualized by co-staining with DAPI.

### Trans-well migration assay

H1975 cells were grown in 6-well plates and treated with siRNAs for Numb and Numb-L for 24 hours. After 48hrs of siRNA knockdown, cell monolayers were harvested, counted and cell concentration was adjusted to 2×10^4^ viable cells/200μl of serum free medium. The cell suspension was seeded on top of a Transwell insert with 0.8 μm pore diameter (Millipore) and placed on a 24 well cell culture plate. At the bottom of the Transwell insert, 10% fetal bovine serum was added as chemo-attractant and plate was incubated at 37°C for 18 hours. Medium was aspirated from assay plate and Transwell inserts. Cells that did not migrate were removed by gently swabbing the inside of each insert using cotton swabs. After 3 washes with PBS, cells were fixed and stained with a 0.5% crystal violet solution for 10 minutes. After the cells have been stained, the inserts were washed thoroughly with water until the water runs clear. Inserts were dried completely before visualizing with a microscope. The image is from 72 hrs experiment just to match the time course of IF experiments.

### Proliferation assay

MTS assay (CellTiter 96 Aqueous One Solution Cell Proliferation Assay, Promega) was performed to assess the cell viability after 72 hours and 96 hours. The experiments were repeated 3 times each with five technical replicates. The error bar represents the % standard deviation of the MTS values normalized with controls. We followed the standard protocol as per the manufacturer's instruction (Promega, CellTiter 96^®^ AQueous One Solution Cell Proliferation Assay).

### Kaplan-Meier plots

All survival analysis plots were generated using ProgGene^13^.

### Mathematical model of the Notch-EMT-Numb axis

The mathematical model of the Notch-EMT-Numb axis describes the dynamics of the molecular species of the EMT regulatory circuit (miR-34, miR-200, Snail, Zeb), the Notch signaling pathway (Notch receptor, Delta, Jagged, NICD) and Numb according to the schematic of Fig. 1B. The temporal dynamics of the species in the circuit is modeled via a system of ordinary differential equations. The complete set of equations is presented in the Supplementary Information, Mathematical Model – Notch-Numb-EMT circuit. Additionally, the section “Implementation of the miR-34 inhibition on Numb” of the SI models the post-translational inhibition of Numb by miR-34. Every chemical species is characterized by its own basal production and degradation rate. Furthermore, the production rate of any species can be modulated by transcriptional/translational regulation. Details on how such interactions are modeled are given in the “Quick guide to equations and assumptions”. All used parameters are discussed in the Supplementary Information, Mathematical Model – Parameters estimation. In the single cell model (Fig. 2), the isolated cell is described by the levels of all proteins, transcription factors (TFs) and micro-RNAs (miRs) in the model, always expressed in number of molecules. Additionally, the cell can be exposed to an external level of Notch receptor and ligands Delta and Jagged *(N*_ext_, *D*_ext_, *J*_ext_) to mimic an incoming signal from neighboring cells. The cell is classified as epithelial (E), hybrid (E/M) or mesenchymal (M) according to its level of micro-RNA 200 (miR-200 <5000 molecules: M; 5000<miR-200<15000: E/M; miR-200>15000: E).

**Figure 1.**
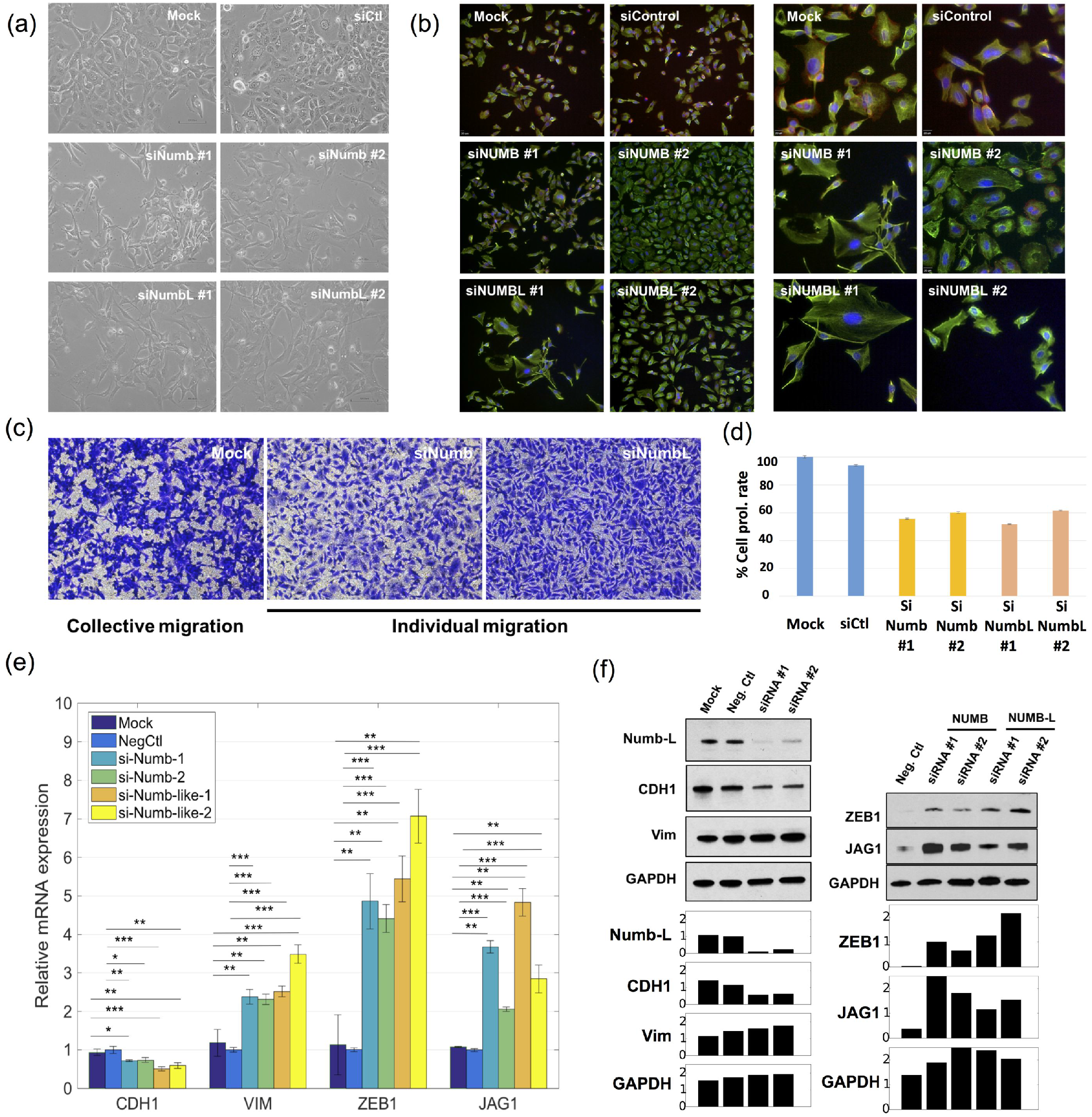
Knockdown of Numb or Numbl induces a full EMT in H1975 cells. (a) Bright-field microscopy for mock H1975 cells, H1975 with control siRNA, and H1975 with siRNA against Numb or Numbl. (b) Immunoflourescence images where red stains for CDH1 (E-cadherin), green for VIM (Vimentin) and blue for DAPI (nucleus). Left panel, magnification 100X, right panel, magnification 200X (c) Transwell migration images for mock H1975 cells, and those treated with siRNA against Numb or Numbl. (d) Effect of Numb- or Numbl-KD on proliferation of H1975 cells. N=5 for each technical replicate. Error bars represent standard error of mean (S.E.M.). (e) RT-PCR measurements of levels of CDH1 (E-cadherin), VIM (vimentin), ZEB1 and JAG1 in cells treated with siRNA either against Numb or Numbl. (f) Western blot measurements for CDH1, VIM, ZEB1 and JAG1 in cells treated with either Numb or Numbl. Left panel represents siRNA against NUMBL. Bar charts show measurement quantification done with the software imageJ. Intensities are normalized over the negative control of Numb-L (left) and the siRNA#1 of ZEB (right), respectively. Corresponding NUMB results are in SI, Fig. S1. Neg Ctľ indicates negative control.

**Figure 2.**
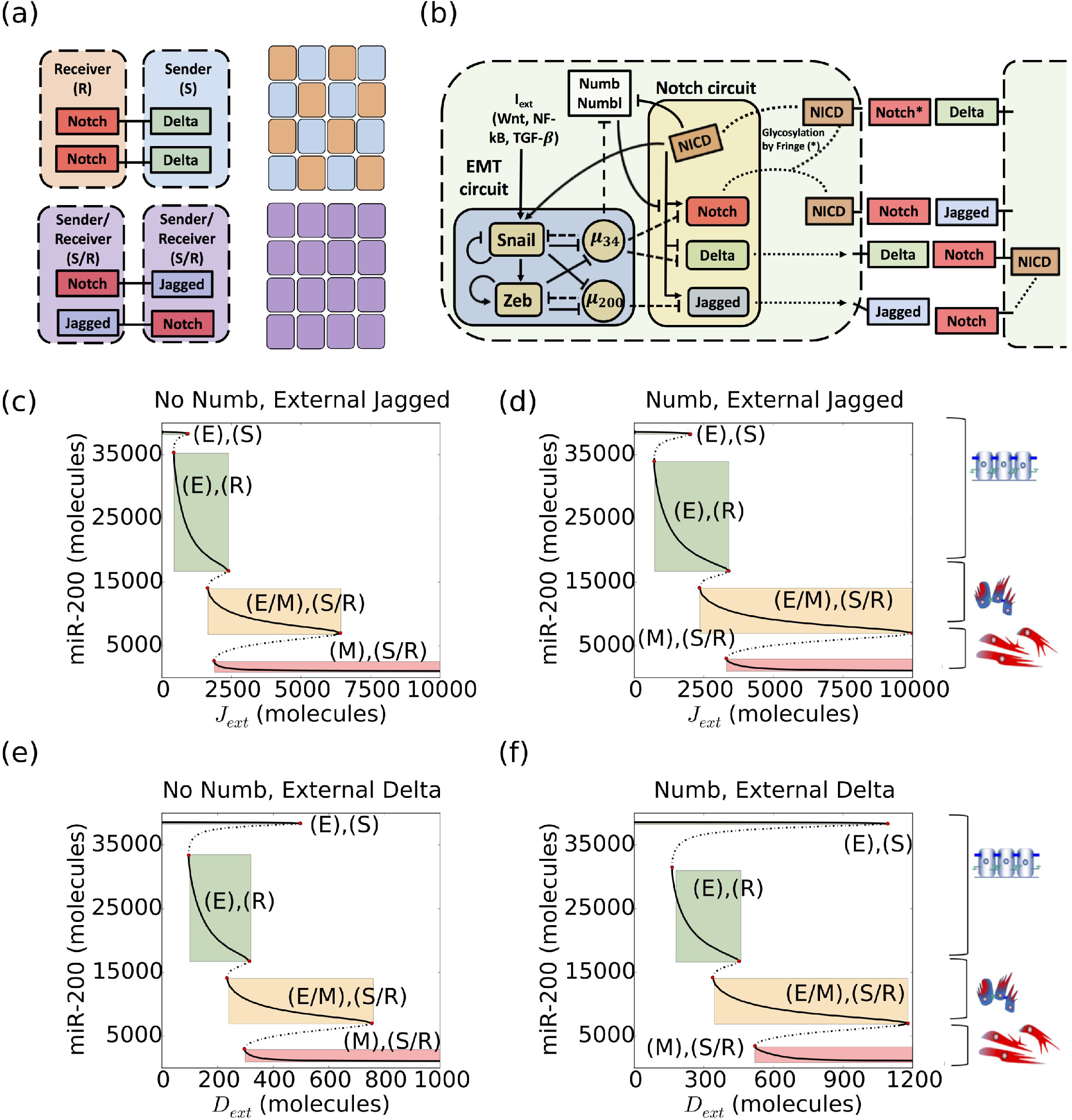
Coupling of Notch signalling with the EMT regulatory circuit and bifurcation curves of miR-200 for Notch-EMT and Numb-Notch-EMT circuits. (a) Cells communicating via Notch-Delta signalling exhibit divergent cell fate. One cell has high levels of Notch and low levels of Delta, thus behaving as a Receiver (R), whereas the other has high Delta and low Notch, thereby behaving as a Sender (S). Conversely, cells that interact through Notch-Jagged signalling assume a similar phenotype. Both cells have a high concentration of Notch and Jagged (Sender/Receiver, S/R). At a multi-cell level, Notch-Delta signalling can generate a ‘salt-and-pepper’ pattern where a sender cell is surrounded by receiver cells and vice-versa. On the other hand, Notch-Jagged signalling generates a uniform distribution of similar S/R cells. (b) Schematics of the connection between the EMT regulatory unit and the Notch signalling circuit. The microRNA miR-34 inhibits Notch and Delta, while miR-200 inhibits Jagged and NICD activates Snail. Numb inhibits Notch while being inhibited by NICD. Additionally, miR-34 inhibits Numb. (c) Bifurcation curve of the level of miR-200 as a function of external Jagged concentration Jext without Numb inhibition acting on Notch. Here, the external concentration of Delta is fixed to zero. Thick and dashed black lines represent stable and unstable steady states, respectively. Cartoons alongside the figure depict which steady states correspond to which EMT phenotypes. Coloured rectangles highlight the interval of stability (parameter on the x-axis) and the corresponding level of the microRNA miR-200 (y-axis) for the different states. (d) Same as (c) in presence of the Numb-related interactions in the system. (e) Bifurcation curve of miR-200 as a function of external Delta concentration Dext without Numb. The external concentration of Jagged is fixed to zero. (f) Same as (e) as Numb is inserted in the system. In all simulations, the concentration of external Notch is fixed to N_ext_=10000 molecules. Bifurcation curves of all proteins and micro-RNAs in the model are shown in Fig. S2-S5.

The multi-cell model (Fig. 3–4–5) extends the single cell model to a two-dimensional layer of 50x50 cells arranged in a hexagonal lattice. In this model, each cell is still described by its own set of proteins, TFs and miRs. Additionally, the Notch receptors and ligands of every cell can bind with the receptors and ligands of any neighboring cell. Also, cells in the layer can be exposed to a constant level of soluble ligand (*D*_sol·_, *J*_sol·_) as in Fig. 5 or to an external EMT-inducer *I*_ext_.

**Figure 3.**
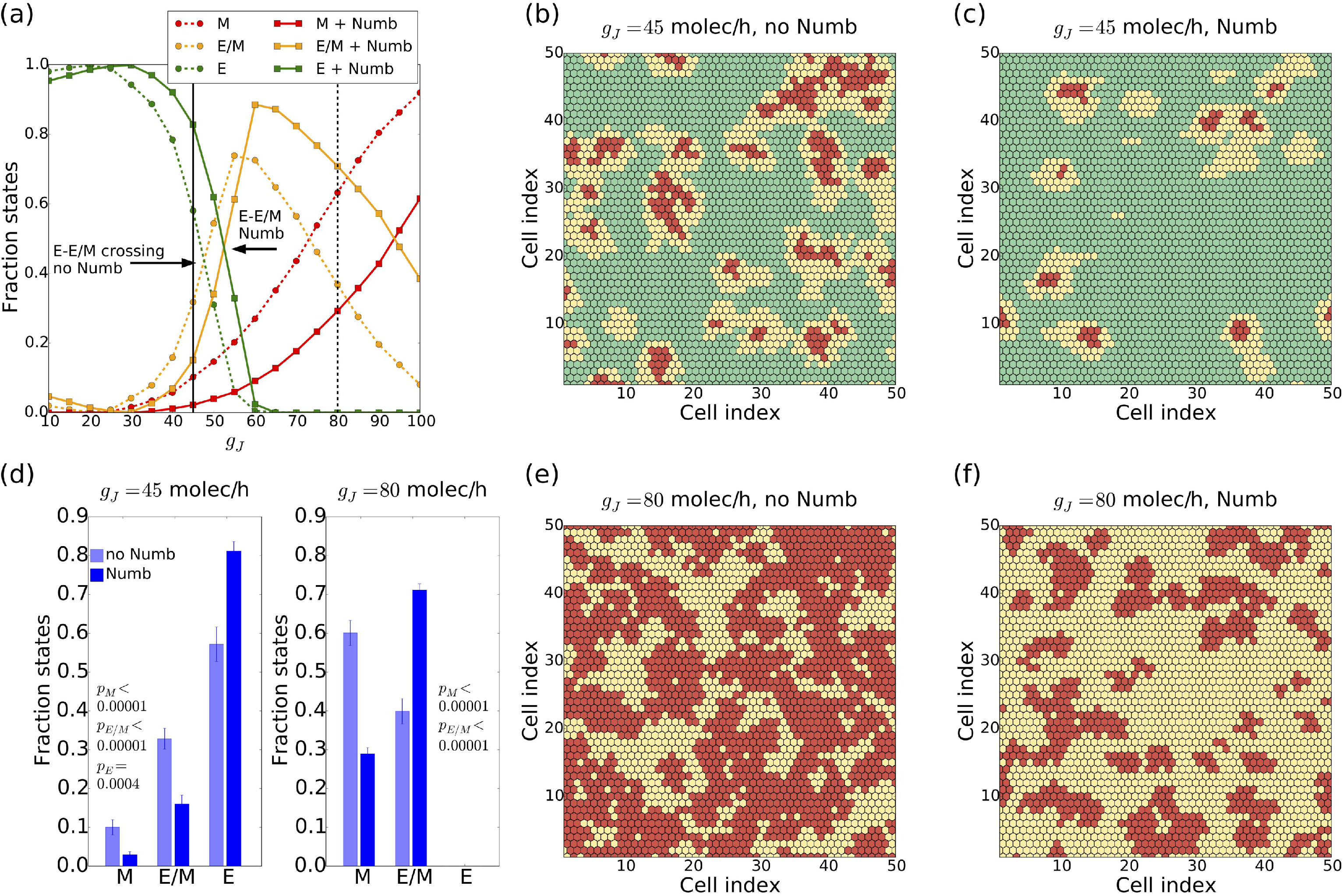
Effect of Numb on tissue patterning for Jagged-dominated Notch signalling. (a) Fraction of epithelial (E), epithelial/mesenchymal (E/M) and mesenchymal (M) cells as a function of the production rate of the ligand Jagged (g_J_) in a 2-dimensional 50x50 layer of cells in the absence or presence of Numb interactions (dashed and continuous lines, respectively). The vertical continuous and dashed black lines depict the values of gj used in (b),(c) and (e),(f), respectively. The required production rate to observe the same fraction of E and E/M cells (green and yellow curves) is increased when Notch is inhibited by Numb. The crossing between hybrid E/M and mesenchymal (yellow and red curves) is shifted toward a larger production rate as well. (b) Snapshot of a 2-dimensional layer of cells interacting without Numb for gJ=45 molecules h^1^ corresponding to the E-E/M crossing without Numb. (c) Same as (b) for the Notch-EMT-Numb circuit. (d) Average fraction of E, E/M and M cells for g_J_=45 molecules h^1^ and g_J_=80 molecules h^1^. For g=45 molecules h¯, Numb decreases the fraction of both hybrid and mesenchymal cells. At g=80 molecules h¯ all cells have undergone partial or complete EMT, but Numb reduces the fraction of mesenchymal cells. Averages are computed over 10 simulations starting from different randomly chosen phenotype distributions. (e) Snapshot for g_J_ =80 molecules h^-^ in absence of Numb. (f) Same as (e) in presence of Numb. The production rate of Delta is fixed at g_D_=20 molecules h^-^ in all plots. The fractions of states and the snapshots were taken after a transient of 120 h starting from the same randomized initial conditions.

**Figure 4.**
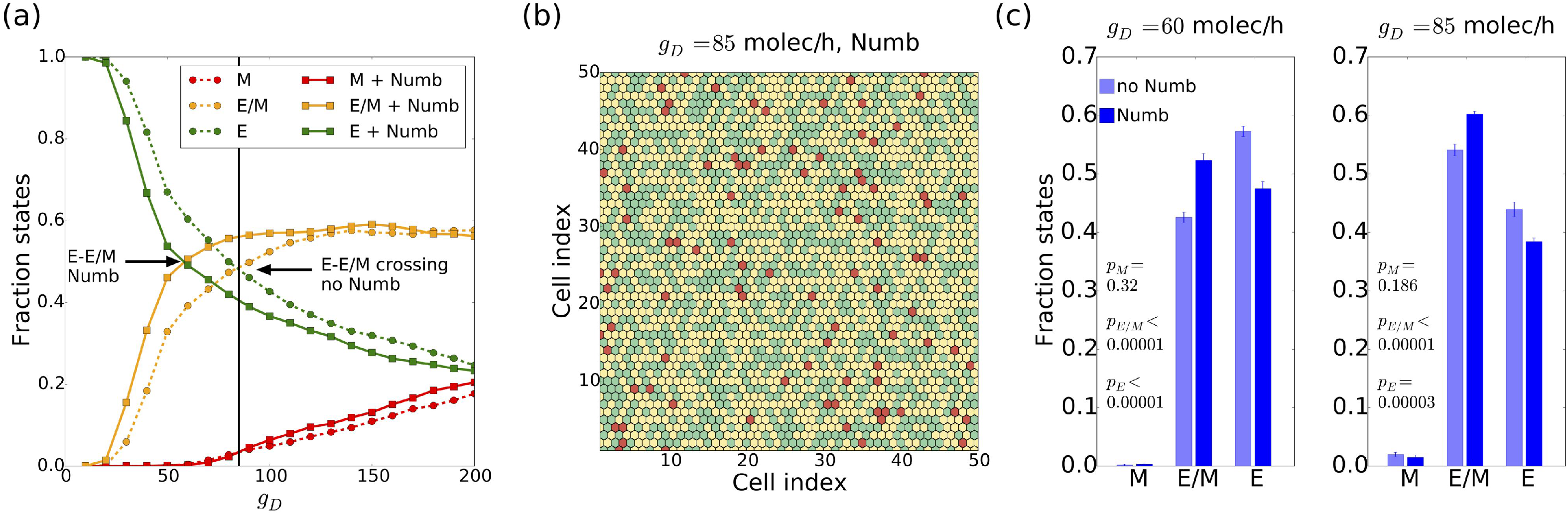
Effect of Numb on tissue patterning for Delta-dominated Notch signalling. (a) Fraction of epithelial (E), epithelial/mesenchymal (E/M), mesenchymal (M) cells as a function of the production rate of the ligand Delta in a 2-dimensional 50x50 layer of cells in the absence or presence of Numb interactions, i.e. Notch-EMT and Notch-EMT-Numb circuits (dashed and continuous lines, respectively). The vertical continuous line depicts the value of gD used in (b) The required production rate to observe a 1:1 ratio of E and E/M cells (crossing of green and yellow trajectories) is decreased when Notch is inhibited by Numb. (b) Snapshot of 2-dimensional cell layer for g_D_=85 molecules h^1^ corresponding to the E-E/M crossing for Notch-EMT-Numb circuit. (c) Average fraction of cells in the three states for two values of gD corresponding to the two crossing points between E and E/M. Averages are computed over 10 simulations starting from different randomly chosen phenotype distributions. The production rate of-1 Jagged is fixed at gJ=20 molecules h^−1^ in all plots. The fractions of states and the snapshots were taken after a transient of 120 h starting from the same randomized initial conditions.

**Figure 5.**
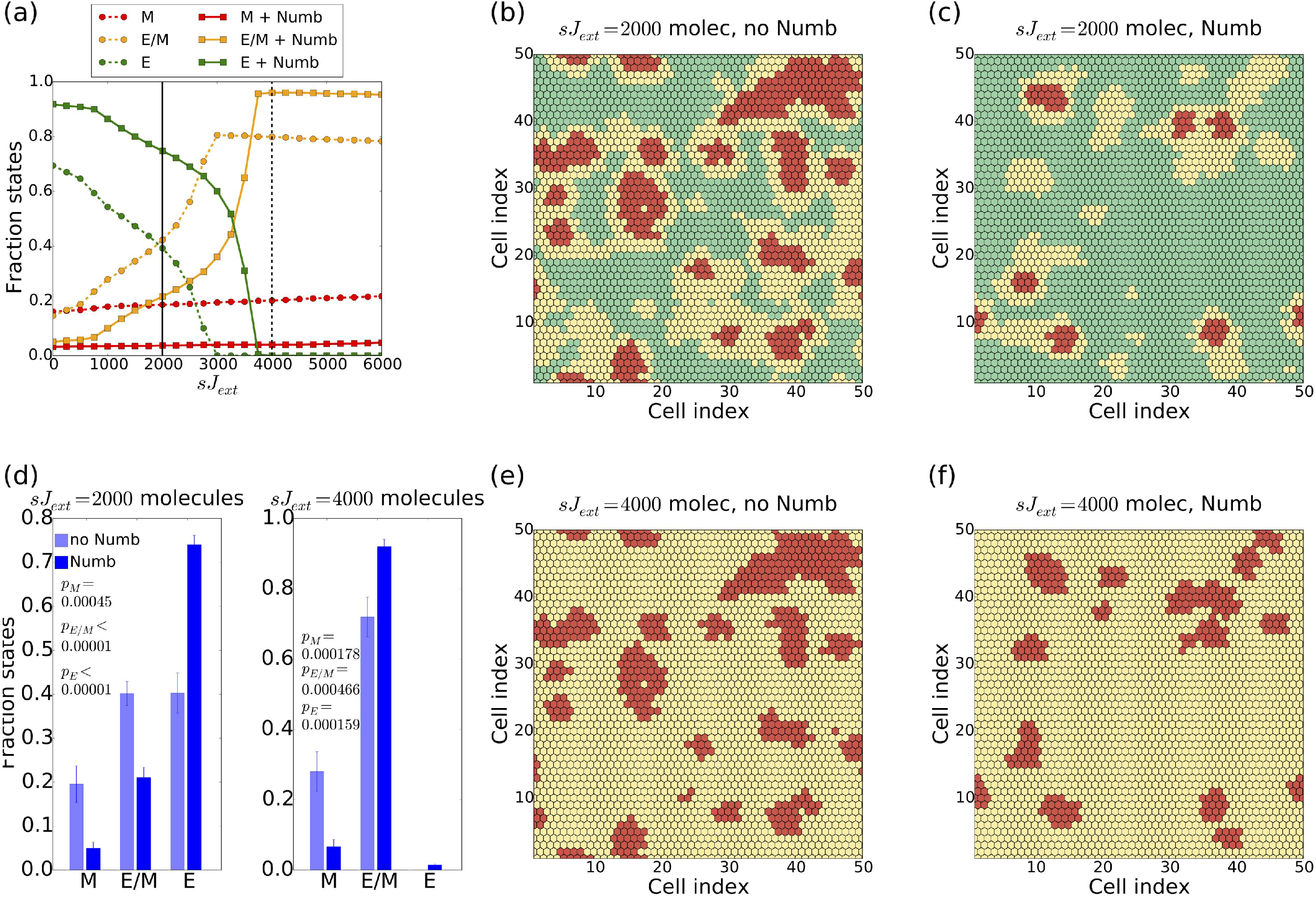
Effect of Numb in presence of soluble Jagged when Notch signalling is Jagged-dominated. (a) Fraction of epithelial (E), epithelial/ mesenchymal (E/M) and mesenchymal (M) cells as a function of the concentration of soluble Jagged sJext in a 2-dimensional 50x50 layer of cells in the absence or presence of Numb interactions (dashed and continuous lines, respectively). Cells in the lattice communicate preferentially through Notch-Jagged signalling (g_J_=45 molecules h¯, g_D_=20 molecules h¯). The vertical continuous and dashed black lines depict the values of sJ_ext_ used in (b),(c) and (e),(f), respectively. (b)-(c) Snapshot of the 2-dimensional cell layer for sJ_ext_=2000 without Numb (b) and with Numb (c): in the presence of Numb the formation of clusters is considerably restricted and the fraction of mesenchymal cells is drastically decreased. (d) Average fraction of E, E/M and M cells for sJ_ext_=2000 and sJ_ext_=4000 molecules. In both cases, Numb strongly diminishes both partial and complete EMT. The averages are computed over 10 simulations starting from different randomly chosen phenotype distributions. (e)-(f) Snapshot of the 2-dimensional cell layer for sJ_ext_=4000 without Numb (e) and with Numb (f): in the presence of Numb the fraction of mesenchymal cells is drastically decreased. Fractions of states and snapshots were measured after a transient of 120 h starting from the configuration of Fig. 4B and 4C as initial conditions for the cases in the absence or presence of Numb, respectively.

### Numerical calculation and plotting

The single cell and the multi-cell systems are implemented and solved numerically using the python numerical library PyDsTool^14^. All plots are realized with the plotting library Matplotlib^15^. All source code is freely available on Github (https://github.com/federicobocci91/Numb_project).

## Results

### Numb knockdown drives hybrid E/M cells to a completely mesenchymal phenotype

As a first step to investigate the effect of Numb on EMT, we knocked down either *Numb* or its homolog Numb-like *(Numbl)* in non-small cell lung cancer (NSCLC) H1975 cells that display a stable hybrid E/M phenotype over many passages *in vitro*.

Knockdown of Numb or Numbl changed the morphology of H1975 cells to being more spindle-shaped, and individual cells stained positive only for mesenchymal marker vimentin (VIM) but not for epithelial marker E-cadherin (CDH1), as compared to the control H1975 cells that coexpress E-cadherin and vimentin stably over many passages^4^ (Fig. 1A, B). Moreover, in transwell migration assays, control H1975 cells exhibited collective cell migration, but *Numb-* or *Numbl-* knockdown H1975 cells displayed individual cell migration (Fig. 1C). These observations mimic earlier observations made in multiple contexts such as mammary gland development^16^, MCF10A cells^17^, MDCK cells^18^, and esophageal cancer cells^19^. Further, knockdown of *Numb* or *Numbl* lead to inhibition of cell proliferation, a trait also typically associated with EMT progression^20^ (Fig. 1D). Similar effect on inhibited proliferation was also observed for knockdown of GRHL2 – another proposed PSF – in lung^4^ and ovarian^21^ cancer cells.

Consistently, *Numb-* or *Numbl-* knockdown increased the mRNA and protein levels of (a) mesenchymal marker Vimentin, (b) EMT-inducing transcription factor ZEB1, and (c) Notch ligand JAG1. Conversely, mRNA and protein levels of E-cadherin were decreased (Fig. 1E-F, S1). Put together, these observations indicate that knockdown of *Numb* or its homolog *Numbl* in stably hybrid E/M cells drive them towards a more mesenchymal phenotype, thereby strongly supporting our hypothesis that Numb or Numbl can independently stabilize a hybrid E/M phenotype and act as a brake on complete EMT progression.

### Numb inhibits a complete EMT at a single-cell level by restricting Notch signalling

To better characterize the effect of Numb on the dynamics of epithelial-hybrid-mesenchymal transitions, we extend our previously defined mathematical model to include Numb regulation of Notch. As mentioned earlier, Notch signalling takes place when Notch (trans-membrane receptor) of one cell binds to Delta or Jagged (transmembrane ligands) of the neighbouring cell(s). Signalling through different ligands, Delta or Jagged, lead to a different phenotypic patterning at a multi-cellular level. Notch-Delta (N-D) signalling between two cells creates divergent cell fates – one cell behaves as a Receiver (high receptor, i.e. Notch, low ligand, i.e. Delta) and the other behaves as a Sender (low receptor, i.e. Notch, high ligand, i.e. Delta). Conversely, Notch-Jagged signalling leads to convergent cell fates – both cells behave as hybrid Sender/Receiver (high receptor, i.e. Notch, high ligand, i.e. Jagged)^10,11^ (Fig. 2A). This trait of the Notch-Jagged signalling can contribute to the formation of clusters of hybrid E/M cells by ‘lateral induction’ of a hybrid E/M phenotype^8^, due to the coupling between Notch and EMT circuits as shown in Fig. 2B, where Notch activates SNAIL, an EMT-inducing transcription factor, and miR-34 and miR-200 families – guardians of an epithelial phenotype^1^ – inhibit Notch, Delta, and Jagged (details of the circuit given in SI section…). EMT can be driven by either Notch-Delta or Notch-Jagged signalling^8^. Numb and Notch form a mutually inhibitory feedback loop^9^ (Fig. 2B). For modelling purposes, we consider Numb and Numbl to be equivalent and group them into one variable.

As a first step to decode how Numb affects EMT at a single-cell level, we compared the intracellular dynamics of coupled Notch-EMT and Notch-EMT-Numb circuits as a function of fixed levels of external ligands – J_ext_ and D_ext_ – that represent the average concentration of Delta and Jagged available at the surface of the neighbouring cells. Previous work has shown that activation of Notch signalling by either Delta or Jagged can induce a partial or complete EMT in epithelial cells^8,22,23^. Consistently, we observed cells attaining a partial or complete EMT in both cases, i.e. with and without Numb (Fig. 2C-F).

In absence of Numb, at a low external concentration of either ligand, a cell maintains its epithelial phenotype and can behave as either a sender or a receiver – (E), (S) or (E), (R). At higher ligand concentrations, the cell transits to a hybrid E/M state and can act both as sender and receiver – (E/M), (S/R). Eventually, at an even higher concentration of ligands, the cell undergoes a complete EMT – (M), (S/R) (Fig. 2C, 2E). A similar trend is observed in the presence of Numb, but the range of existence of these different states is altered. Numb enlarges the range of J_ext_ and D_ext_ values for which the (E), (R) and (E), (S) state exist (compare the width of the green rectangle in Fig. 2D vs. that in Fig. 2C, and in Fig. 2F vs. that in Fig. 2E). Furthermore, the range of values of external ligand concentrations for which the cell maintains a stable hybrid E/M state – (E/M), (S/R) – is increased (compare the width of orange rectangle in Fig. 2D vs. that in Fig. 2C, and in Fig. 2F vs. that in Fig. 2E). Consequently, cells can maintain a (E/M), (S/R) state at much higher levels of external ligands. Thus, a transition towards a complete EMT state is inhibited. In other words, cells need a stronger stimulus to attain a mesenchymal state (compare the value of J_ext_ at the left end of red rectangle in Fig. 2D vs. that in Fig. 2C, and the value of D_ext_ at the left end of red rectangle in Fig. 2F vs. that in Fig. 2E). Altogether, these results indicate that Numb can restrict the progression of a complete EMT, and may stabilize both epithelial and hybrid E/M phenotypes at a single-cell level.

To probe the robustness of these results, we conducted a sensitivity analysis by assessing the change in the level of NICD resulting from a small variation of the model’s parameters. Our results are robust upon parameter variation (Fig. S6), albeit a higher sensitivity was observed for the inhibition of Notch by Numb than *vice-versa* (Fig. S7).

Overall, our results suggest that Numb or Numbl can act as a PSF that can stabilize a hybrid E/M phenotype at a single-cell level.

### Numb alters the composition of clusters of non-epithelial cells at a tissue level

After evaluating the effect of Numb on EMT at a single-cell level, we compared the dynamics of Notch-EMT and Notch-EMT-Numb circuits at a tissue level by simulating a two-dimensional lattice of 50x50 cancer cells communicating with one another via Notch signalling. Specifically, we studied the relative abundance of epithelial (E), hybrid (E/M) and mesenchymal (M) cells and the spatial patterns that these subpopulations form in this lattice, at different production rates of Jagged (g_j_) and Delta (g_D_), starting from random initial conditions.

We first compared the tissue-level dynamics of Notch-EMT and Notch-EMT-Numb circuits, when cells mainly interact via Notch-Jagged signalling (Fig. 3). Notch-Jagged signalling can promote the formation of clusters containing hybrid E/M and M cells^8^. At low levels of Jagged production (g_J_=45 molecules/h), Notch-Jagged signalling is only weakly activated. In this regime, additional inhibition on this signalling brought by Numb decreases the abundance of both hybrid E/M and mesenchymal cells (Fig. 3A, solid vertical black line), thus halting EMT progression. Consequently, Numb reduces the frequency of clusters containing hybrid E/M and M cells (compare Fig. 3C with Fig. 3B). To quantify the changes induced by Numb, we counted the fraction of epithelial, hybrid and mesenchymal cells over many different simulations (each simulation has slightly different initial conditions). For g_J_=45 molecules/h, Numb significantly reduces the number of cells in a partial or complete EMT state, and consequently increased those in an epithelial state (Fig. 3D, left).

When comparing Notch-EMT and Notch-EMT-Numb circuits for higher production rates of Jagged (g_J_=80 molecules/h), a different role of Numb is revealed. In this regime, a strong activation of Notch-Jagged signalling is capable of pushing most cells to either a partial or a complete EMT (Fig. 3A, dashed vertical black line). However, Numb inhibits the accumulation of cells in a complete EMT state while increases those in a hybrid E/M state (compare Fig. 4F with Fig. 3E). This behaviour of Numb as a PSF is reminiscent of its role seen both in H1975 cells (Fig. 1) and in our single-cell simulations (Fig. 2). This effect of Numb has been quantified by measuring the change in the fraction of hybrid E/M versus mesenchymal cells in the absence or presence of Numb (Fig. 3D, right) for the case of a large production rate of Jagged that can push ~75% cells in a complete EMT state (g_J_=80 molecules/h).

Finally, to quantify the spatial co-localization of hybrid E/M cells, we counted how many cells adjacent to a hybrid E/M cell exhibited the same, i.e. a hybrid E/M, phenotype (Fig. S8). For the case of weakly activated Notch-Jagged signalling corresponding to lower gJ (Fig. 3B-C), the average number of hybrid E/M neighbours for a hybrid E/M cell decreased (compare Fig. S8 middle panel with S8 left panel) due to a decreased total frequency of hybrid E/M cells. However, an increased production of Jagged (Fig. 3E-F) can counteract this effect of Numb and consistent with previous reports^8^, it can significantly increase the co-localization of hybrid E/M cells (Fig. S8, right).

Similar to the Notch-Jagged case, we compared the tissue-level spatiotemporal dynamics for Notch-EMT and Notch-EMT-Numb circuits in a lattice of cells that communicate with one another predominantly via Notch-Delta, instead of Notch-Jagged (Fig. 4). Inhibition of Notch signalling by Numb reduces NICD levels^9^, thereby effectively relieving the inhibition of Delta by NICD. This effective increase in the levels of Delta can potentiate Notch signalling in neighbouring cells and thus promote EMT in those cells. As a result, in the case of Notch-EMT-Numb circuit and Delta-dominated signalling, lower basal production levels of Delta (g_D_) can enable transitions into the hybrid E/M state, as compared to that required to observe these transitions in the absence of Numb (compare the solid yellow curve with dotted yellow curve in Fig. 4A). Therefore, at a fixed production rate of Delta (g_D_), the Notch-EMT-Numb circuit can induce significantly more epithelial cells to attain a hybrid E/M phenotype as compared to that by Notch-EMT circuit (Fig. 4C). Contrary to the case of strong Notch-Jagged signalling case, here the increase of the hybrid E/M cell population is mostly due to a decrease in the frequency of epithelial cells. Despite the effect of Numb in altering the ratio of cells in a hybrid E/M and epithelial phenotype, it did not alter the predominant ‘salt-and-pepper’ pattern of epithelial and hybrid E/M cells (Fig. 4B, S9). Such pattern formation is a cornerstone of Notch-Delta signalling as observed in multiple biological contexts^10^.

Collectively, these results suggest that irrespective of the ligand activating Notch signalling – Delta or Jagged – Numb can increase the number of cells in a hybrid E/M phenotype at both a single-cell and a tissue-level.

After investigating the effect of Numb on Notch-EMT circuit, both for Jagged-dominated and Delta-dominated scenarios, we explored the effect of Numb on modulating the paracrine version of Notch signalling, i.e. when cells are exposed to soluble Delta (sD_ext_) or soluble Jagged (sJ_ext_), in addition to membrane-bound ligands (juxtacrine signalling) considered so far in our simulations. Consistent with our results, Numb reduced the frequency of cells in a mesenchymal phenotype in a cohort of cells that were exposed to either soluble ligand (Fig. 5A-D, S10-12). Similar to previous observations (Fig. 3), an increase in soluble Jagged concentration rescues the cluster frequency, but these clusters predominantly contain hybrid E/M cells and not mesenchymal cells (Fig. 5E-F, S12). These effects of Numb on paracrine signalling are more prominent in case of Jagged-dominated juxtacrine signalling instead of Delta-dominated juxtacrine signalling (Fig. S10, S12).

In addition, the presence of soluble Jagged in the microenvironment has a crucial consequence on the dynamics of cell fractions in different phenotypes. It can increase the lifetime of transiently observed clusters of hybrid E/M and mesenchymal cells for both Delta-dominated and Jagged-dominated juxtacrine signalling. Without the presence of soluble Jagged, as the Notch-EMT system tends towards a stable equilibrium, hybrid E/M and epithelial cells arrange themselves in a ‘salt-and-pepper’ pattern for Delta-dominated signalling. On the other hand, in the case of Jagged-dominated signalling, cells in hybrid E/M and M phenotypes tend to an epithelial switch (Fig. S13A, B). The presence of external soluble Jagged stabilizes the hybrid E/M phenotype, thereby further increasing the lifetime of the clusters in the Notch-Jagged signalling case (Fig. S13C, D).This effect of soluble Jagged in the extracellular environment may help explain how soluble Jagged can drive the cells towards a cancer stem cell phenotype^24^ which is often correlated with a hybrid E/M phenotype^1^.

It should be noted that soluble Delta or Jagged driven signalling is fundamentally different from the formation of intercellular feedback loops between Notch-Delta or Notch-Jagged signalling that are responsible for different patterns formed in Delta-dominated and Jagged-dominated signalling. When soluble ligands – whether Jagged or Delta – activate Notch signalling, the cells only behave as ‘receiver’ or ‘target’ in case of either ligand, without any tangible feedback on the amount of these soluble ligands. Therefore, Numb similarly affects the dynamics of the system in case of soluble Delta or soluble Jagged driven signalling.

### External EMT induction can overcome the inhibition of EMT by Numb

We next considered the effect of an external EMT inducer such as TGF-β (I_ext_) that activates SNAIL. As shown in the case of Jagged-dominated Notch signalling, high levels of I_ext_ significantly increase the number of cells in a fully mesenchymal phenotype (Fig. S14A). Consistent with our single-cell results, Numb acts as a molecular brake on EMT (compare the dotted curves against solid curves in Fig. S14A), and therefore stronger induction of EMT is needed to increase the number of mesenchymal cells. Intriguingly, the frequency of cells in a hybrid E/M phenotype in this case is minimal (Fig. S14B-D). These results may help explain why ectopic overexpression of ligand-of-Numb X (LNX) – an ubiquitin ligase that targets Numb for degradation – can enhance TGF-β induced EMT^25^.

Conversely, when cells communicate predominantly via Notch-Delta signalling, Numb can mildly assist EMT induction and increase the fraction of mesenchymal cells in the population (Fig. S15 A-D). This differential effect of Numb in regulating Notch-Jagged and Notch-Delta signalling is further confirmed by assessing the temporal changes in fraction of epithelial, hybrid E/M and mesenchymal cells (Fig. S13 E-F), where Numb slows down the induction of EMT for Notch-Jagged signalling but enhances that for Notch-Delta signalling.

Lastly, reproducing the experimental setup of Fig. 1, we set up a simulation where all cells in the layer are initially hybrid E/M, and compare the dynamics of the Notch-EMT vs. Numb-Notch-EMT circuit in presence of EMT-induction. Confirming the experimental observation, cells that lack Numb (Notch-EMT case) becomes mesenchymal on a timescale of 4-5 days (Fig. S16).

Finally, we considered the effect of another recently reported feedback regulation in coupled EMT-Notch circuit – the relatively weak inhibition of Numb by miR-34^26^. Owing to its weak strength, miR-34 only subtly alters the effect of Numb on EMT and Notch signalling (Fig. S17- S21).

### Higher *Numb* or *Numbl* levels predict poor patient survival

The ability of Numb to stabilize a hybrid E/M phenotype and increase the number of hybrid E/M cells in CTC clusters strongly suggested its potential role as a PSF. These observations are conceptually similar to previous computational and experimental attempts in identifying PSFs such as OVOL2, GRHL2, and ΔNp63α that can induce and/or maintain a hybrid E/M phenotype, as observed in breast cancer and NSCLC cells^4,27,28,7,29^. Intriguingly, high levels of such PSFs have been proposed to predict poor survival, for instance, GRHL2 is associated with poor survival across different breast cancer subtypes^30^, and high ΔNp63α correlate with poor survival in ER^-^/HER2^-^ patients^27^. Thus, we next investigated the association of *Numb* or *Numbl* with patient survival.

High levels of *Numb* or *Numbl* were found to associate with poor overall survival (OS) and relapse-free survival (RFS) in multiple independent lung cancer datasets (Fig. 6, top and middle rows) as well as in ovarian cancer datasets (Fig. 6, bottom row). Our results are consistent with the reported association of high levels of Numb with poor survival in hepatocellular carcinoma^31^, with tumour recurrence and poor overall post-operative survival in esophageal squamous cell carcinoma^32^, and that of with poor overall survival in NSCLC patients^33^. Also, siRNA against *Numbl* inhibited the ability of lung cancer cells to form liver metastasis^33^, thus reinforcing a role of Numb and/or Numbl in driving tumor aggressiveness.

**Figure 6.**
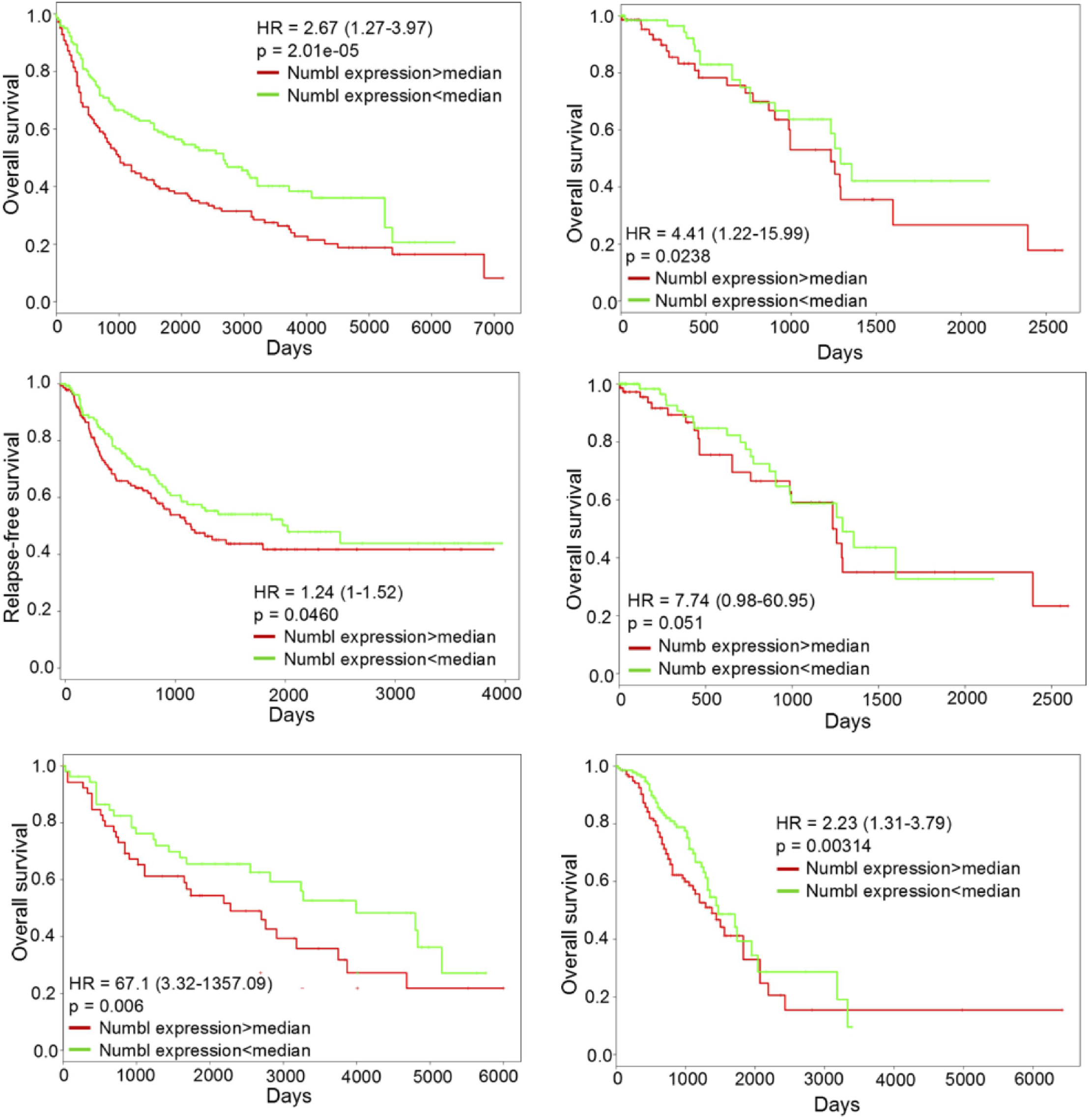
Numb or Numbl can predict poor survival. Survival plots for GSE30219 (n=282), GSE 41271 (n=275), GSE 73614 (n=106) (left panel, top-bottom), TCGA-LUAD (n=150), TCGA-LUAD (n=150), GSE9891 (n=276) (right panel, top-bottom).

Low levels of Numb and/or Numbl, indicative of cells that have completely progressed to a mesenchymal phenotype, associate with a better survival, thereby reinforcing the emerging notion that a partial EMT, instead of a full EMT, may be a better marker for tumour aggressiveness^1^. These notions are supported by recent clinical evidence indicating that singlecell migration (a canonical readout of full EMT) happens extremely rarely, if any, in cancer dissemination^34^.

## Discussion

Notch signalling pathway is implicated in multiple hallmarks of cancer including metastasis and angiogenesis, and other clinically insuperable aspects such as drug resistance^35^. Here, we investigate how Numb – an inhibitor of Notch signalling – modulates EMT, a process that can contribute to both metastasis and drug resistance. Our results suggest that Numb can prevent cells from undergoing a complete EMT by modulating Notch signalling, and knockdown of Numb can induce a full EMT in H1975 cells exhibiting a stable hybrid E/M phenotype. These observations resonate well with recent observations that knockdown of Numb can induce EMT in MCF10A cells^17^, as well as during mammary gland development^16^. Consistently, Numb overexpression led to a loss of mesenchymal markers and features, thereby pushing the cells to an epithelial state^16,17^. Collectively, these results suggest that Numb can act as a PSF for a hybrid E/M phenotype by altering Notch signalling. Its role as a PSF is bolstered by its association with poor patient survival, a trait previously noted for other PSFs such as OVOL2 and GRHL2.

Knockdown of Numb can not only drive a complete EMT, but also stunt the ability of NSCLC cells to form liver metastases^33^. This effect of Numb corroborates its observed role in predicting poor survival across cancer types, and reinforces strongly the emerging notion that a hybrid E/M phenotype instead of a full EMT may be the hallmark of tumour aggressiveness^1,36,37^.

The effect of Numb on tissue-level patterning is reminiscent of glycosyltransferase Fringe that can alter the binding affinity of Notch with Delta and Jagged, thus affecting tissue patterning in a layer of cells^11^. Both Numb and Fringe tend to antagonize Notch-Jagged signalling predominantly (Fig. S22-S23). This selective inhibition of Notch-Jagged signalling – an axis involved in drug resistance and colonization^8,38,39^ – may help rationalize, at least in part, multiple experimental observations, such as (a) Numb and/or Fringe is/are often lost in many cancer types, including aggressive ones such as basal-like breast cancer (BLBC)^40-42^, (b) *Numbl* knockdown increases chemoresistance and tumorigenic properties in cell lines of different origins – HeLa (cervix), T47D (breast) and AX (sarcoma)^43^, and (c) lunatic fringe *(Lfng)* suppresses *in vitro* tumorsphere formation in prostate cancer DU145 cells^40^.

Importantly, Notch pathway need not be the sole pathway through which Numb modulates EMT. Numb can directly interact with E-cadherin and regulate its membrane localization, as well as control its endocytosis to retain apico-basal polarity in epithelial cells^44,45^. Knockdown of Numb alters E-cadherin localization and polarity complexes such as Par3, and as a result, decreases cell-cell adhesion and increase cell migration^44^. Besides, Numb can stabilize p53^42^ that can activate family members of miR-200 and miR-34 that can restrict EMT and even drive MET^1^. All these aspects of Numb can be integrated with existing theoretical frameworks to better characterize how Numb affects EMT/MET as well as other traits associated with EMT/MET – immune evasion^46^, tumour-initiation potential^1,36^, and drug resistance^8,47^.

Numb has been specifically well-studied in the context of asymmetric stem cell division both for developmental stem cell lineages ^48^ and cancer stem cells (CSCs)^26^. Therefore, future modelling efforts will benefit from integrating the signalling aspects of Numb with population-level models of stem cell division. Such multi-scale spatial models can be critical in explaining the observed spatial patterns of different subpopulations of CSCs in breast cancer tissues^49^.

To conclude, we found that Numb can help in maintaining hybrid E/M phenotype and prevent a full transition to a mesenchymal phenotype, and its knockdown can release the brake for full EMT. Our theoretical framework offers a platform to assess the role of many players that can regulate cellular plasticity in both cell-autonomous and non-cell-autonomous manner, and proposes another target that may potentially break the clusters of tumour cells in a hybrid E/M phenotype – the key drivers of cancer metastasis^1,3^.

